# SRPK3 regulates back-splicing of *pre-MYOCD* through Phosphorylating SRSF10 to modulate cardiac hypertrophy

**DOI:** 10.64898/2026.06.10.731506

**Authors:** Xiaowei Song, Rui Cai, Wenxia He, Tianhong Ai, Shuting Hu, Jinchao Song, Song-Hua Li

## Abstract

Cardiac hypertrophy represents a hallmark of pathological remodeling in heart failure, involving complex molecular networks that remain incompletely defined. While post-transcriptional regulation is increasingly recognized as a key control layer, the role of splicing kinases in this process requires further elucidation. This study identifies serine/arginine-rich protein kinase 3 (SRPK3) as a central regulator of myocardial hypertrophy through its control of circular RNA (circRNA) biogenesis. We observed marked SRPK3 upregulation in phenylephrine/angiotensin II-treated cardiomyocytes, transverse aortic constriction (TAC) rats, and human heart failure datasets. Gain- and loss-of-function studies demonstrated that SRPK3 overexpression exacerbates hypertrophic phenotypes, including increased cell surface area and upregulation of markers (*ANP*, *BNP*, *Myh7*). Conversely, siRNA knockdown or pharmacological inhibition with the pan-SRPK inhibitor MSC-1186 effectively attenuated these pathological features. Mechanistically, SRPK3 interacts with and phosphorylates the splicing factor SRSF10 at specific serine residues (S129, S131, and S133). This phosphorylation event inhibits the back-splicing of *pre-MYOCD*, resulting in reduced cardioprotective *circ-MYOCD* levels and a concomitant increase in linear MYOCD expression. The depletion of *circ-MYOCD* relieves its competitive inhibition on linear splicing, driving the transcriptional activation of pro-hypertrophic signaling pathways. Crucially, restoring *circ-MYOCD* expression reversed SRPK3-driven pathology, confirming its protective role. These findings establish the SRPK3-SRSF10-*circ-MYOCD* axis as a critical regulatory circuit in cardiac remodeling. By bridging kinase activity with splicing dynamics, this study highlights RNA processing as a vital therapeutic target. Targeting this axis via SRPK3 inhibition or *circ-MYOCD* delivery represents a promising dual-modality strategy for treating heart failure.

## Introduction

Heart failure has emerged as the fastest-growing cardiovascular disease worldwide, imposing a substantial burden on global health systems. Predominantly triggered by chronic hypertension, congenital cardiovascular abnormalities, and myocardial infarction [1, 2], the disease is characterized by an early stage of compensatory cardiac hypertrophy. Initially an adaptive response involving cardiomyocyte enlargement and ventricular wall thickening to transiently maintain cardiac output, sustained hypertrophic remodeling eventually progresses to pathological decompensation [3]. This transition impairs both systolic and diastolic functions, significantly elevating the risks of heart failure and sudden cardiac death. Although pathological hypertrophy stems primarily from hemodynamic overload and cardiomyocyte injury, the multifactorial etiology, specifically the dysregulated signaling networks and transcriptional reprogramming driving this process, remains incompletely resolved.

In recent years, post-transcriptional regulation, particularly alternative splicing and RNA processing, has gained recognition as a pivotal layer of control in cardiac hypertrophy. Central to this process are the serine/threonine protein kinases (SRPKs), a specialized family that regulates SR (serine/arginine-rich) splicing factors through phosphorylation [4]. SR proteins are essential for constitutive and alternative pre-mRNA splicing, relying on SRPK-mediated phosphorylation for proper spliceosome assembly and subcellular localization [5]. Beyond splicing, SRPKs influence broader cellular processes, including mRNA maturation, angiogenesis, and cell cycle progression [6]. Among the SRPK isoforms, SRPK3 is uniquely enriched in striated muscle during myocardial development and homeostasis [7–9]. However, its mechanistic role in cardiovascular pathophysiology remains poorly defined, representing a critical gap in our understanding of how splicing dysregulation contributes to heart disease.

Parallel to these findings, circular RNAs (*circRNAs*) have emerged as critical regulators in cardiovascular pathophysiology. These covalently closed non-coding RNAs, which lack 5’ caps and 3’ poly(A) tails, possess exonuclease resistance and superior stability compared to linear RNAs [10, 11]. Generated through the back-splicing of pre-mRNA, *circRNA* biogenesis is orchestrated by canonical spliceosomes and RNA-binding proteins (RBPs), such as QKI and HnRNPA1, which facilitate splice site bridging [12]. Functionally, *circRNAs* regulate cellular processes by acting as microRNA sponges, RBP scaffolds, or direct effectors of cytoskeletal dynamics. In cardiovascular diseases, *circRNAs* exhibit stage-specific dysregulation, modulating myocardial hypertrophy, infarct remodeling, and plaque destabilization [13–15]. For instance, *circSlc8a1* plays context-dependent roles: its suppression via RNA interference or antisense RNA mitigates pressure overload-induced cardiac hypertrophy, whereas its overexpression exacerbates heart failure [16–18]. Similarly, *circCacna1c* has been shown to inhibit oxidative stress-induced necroptosis, highlighting the diverse mechanisms through which *circRNAs* exert their effects [19, 20].

Despite these advances, a fundamental question remains unanswered: how are circRNA biogenesis and splicing networks mechanistically linked during pathological stress? The back-splicing of pre-mRNA into *circRNAs* is tightly regulated by *trans*-acting factors, particularly Serine/Arginine-rich (SR) proteins, whose activity is governed by specific kinases. Given that SRPK3 dictates the phosphorylation state of splicing factors, we hypothesized that it plays a central role in regulating circRNA production under hemodynamic stress. In this study, we bridge this critical gap by identifying SRPK3 as a pivotal regulator of cardiac hypertrophy. We demonstrate that SRPK3 is upregulated in failing human hearts and hypertrophic cardiomyocytes. Mechanistically, we reveal that SRPK3 phosphorylates the splicing factor SRSF10, thereby inhibiting the back-splicing of *circ-MYOCD*. This dysregulation leads to an accumulation of linear *MYOCD* and the activation of downstream pro-hypertrophic signaling. Our findings establish the SRPK3-SRSF10-*circ-MYOCD* axis as a novel therapeutic target, positioning RNA splicing and circRNA biogenesis as promising avenues for intervention in heart failure.

## Materials and Methods

### Human Heart Tissue Collection

Human myocardial tissue samples were procured in strict adherence to the ethical principles outlined in the Declaration of Helsinki. The study protocol was approved by the Institutional Review Board of Naval Medical University (Protocol No. NMU8217021385). Left ventricular specimens were obtained from patients with end-stage heart failure, including those diagnosed with hypertrophic cardiomyopathy (HCM), as well as from non-failing (NF) controls. Tissues were harvested during orthotopic heart transplantation procedures performed at Changhai Hospital between 2012 and 2018. Written informed consent was secured from all participants or their legal representatives prior to sample collection. To preserve RNA and protein integrity, explanted heart tissues were immediately snap-frozen in liquid nitrogen within 2 minutes of surgical resection and subsequently stored at -80°C.

### Euthanasia

For surgical procedures, mice were anesthetized using isoflurane (a halogenated ether). Anesthesia was induced by placing mice in an induction chamber with a gas flow rate of 0.5-1.5 L/min and 3-5% isoflurane until loss of righting reflex was achieved. Following induction, mice were rapidly transferred to a nose-cone delivery system where anesthesia was maintained with a reduced isoflurane concentration (1-2.5%) for the duration of the procedure. Body temperature was supported throughout the entire anesthetic period using a 37°C warming pad. Prior to sacrifice, mice were re-anesthetized with 3-5% isoflurane delivered via the induction chamber. For cardiomyocyte isolation, Sprague-Dawley (SD) rats was anesthetized with a isoflurane concentration (1-2.5%) for 1 minute before sacrificed.

### Mouse Transverse Aortic Constriction (TAC)

Male C57BL/6 mice were anesthetized via inhalation of 1.5-2% isoflurane. Following induction of anesthesia, a left thoracotomy was performed to expose the aortic arch. A 6-0 polypropylene suture was passed beneath the transverse aorta, and a 27-gauge needle was placed parallel to the vessel to serve as a constriction spacer. The suture was then tightened around both the aorta and the needle, and secured with three surgical knots. Immediately after ligation, the needle was carefully withdrawn, resulting in a standardized aortic constriction with a final diameter of approximately 0.4 mm. Sham-operated control mice underwent the same surgical procedure, with the exception of the aortic ligation.

### Isolation and Culture of Cardiomyocytes

Hearts were harvested from 2- to 3-day-old Sprague-Dawley (SD) rats. The ventricles were minced into approximately 1 mm³ fragments and digested with 1% type I collagenase overnight at 4°C. Following centrifugation at 1,500 rpm for 10 minutes, the cells were resuspended in complete DMEM and pre-plated on a 10 cm dish for 2 hours in a 37°C, 5% CO₂ incubator to remove non-myocytes. The enriched cardiomyocytes were then seeded onto 6-well plates and cultured for 36 hours. Subsequently, the medium was replaced with complete DMEM containing 0.1 mM 5-bromo-2′-deoxyuridine (BrdU) for an additional 36 hours to inhibit fibroblast proliferation. Finally, cells were maintained in serum-free DMEM for 24 hours prior to treatment with 100 μM phenylephrine or 5 μM angiotensin II for 48 hours to induce hypertrophy.

### Immunofluorescence Staining of Cardiomyocytes

Primary cardiomyocytes were plated onto coverslips and transfected with either *circ-MYOCD* siRNA (si-*circ-MYOCD*) or control siRNA (CTRL siRNA), or infected with adenovirus overexpressing *circ-MYOCD* (Ad-*circ-MYOCD*) or control adenovirus (Ad-GFP). Forty-eight hours post-transfection/infection, cells were fixed with 4% paraformaldehyde for 30 minutes, permeabilized with 0.1% Triton X-100 for 15 minutes, and blocked with goat serum for 30 minutes. Subsequently, cells were incubated with an α-actinin antibody overnight at 4°C. Following primary antibody incubation, cells were incubated with a Cy3-labeled goat anti-rabbit IgG secondary antibody for 1 hour at room temperature, and nuclei were counterstained with DAPI (100 ng/mL) for 2 minutes. Fluorescence imaging was performed using a laser confocal microscope.

### Construction of *Circ-MYOCD*, SRPK3, SRSF10 Overexpression Adenovirus

The *circ-MYOCD* overexpression vector was constructed by cloning its precursor sequence into the pAdtrack-circRNA shuttle plasmid. Similarly, SRPK3 and SRSF10 overexpression vectors were generated by cloning their respective coding sequences (CDS) into the pAdtrack shuttle plasmid. Following cloning, the plasmids were linearized via PmeI restriction digestion, purified by agarose gel extraction, and transformed into BJ5183 competent cells for homologous recombination with the pAdeasy-1 adenoviral backbone. Recombinant pAdeasy plasmids were isolated and transformed into E. coli DH5α competent cells for amplification. After linearization with PacI, the plasmids were transfected into 293A cells for viral packaging. Cells were harvested 10 days post-transfection, subjected to three freeze-thaw cycles, and centrifuged at 3,000 rpm for 5 minutes. The supernatant containing the adenovirus was collected and used to infect 293 cells for viral amplification. Viral titers were determined by serial dilution assays in 293T cells, and green fluorescence was quantified using fluorescence microscopy.

### Transfection of siRNA into Primary Cardiomyocytes

Three small interfering RNAs (siRNAs) targeting *circ-MYOCD*, *SRPK3*, and other genes, along with a negative control siRNA, were synthesized by Guangzhou Ribo Biotechnology Co. Transfection was performed using Lipofectamine 3000 according to the manufacturer’s instructions.

### RNA Extraction, Reverse Transcription, and Real-Time PCR

Following three washes with PBS, cells were lysed using TRIzol reagent for RNA extraction. Complementary DNA (cDNA) was synthesized from 1 μg of total RNA using HiScript RT III reverse transcriptase. Target RNA expression was quantified by real-time PCR using SYBR Green (TOYOBO) on a Roche LightCycler 480 system.

### Western Blot

Protein samples were denatured by boiling at 95°C for 5 minutes. The denatured proteins were then separated by sodium dodecyl sulfate-polyacrylamide gel electrophoresis (SDS-PAGE). Following electrophoresis, the separated proteins were transferred onto a polyvinylidene fluoride (PVDF) membrane. Subsequently, the membranes were blocked with 5% non-fat milk prepared in Tris-buffered saline containing 0.1% Tween-20 (TBST) for 2 hours at room temperature. After blocking, the membrane was carefully sliced into strips according to the molecular weights of the target proteins. These membrane strips were then incubated with specific primary antibodies overnight at 4°C. Following primary antibody incubation, the strips were washed and incubated with horseradish peroxidase (HRP)-conjugated secondary antibodies for 2 hours at room temperature. After incubation with secondary antibodies, the membrane strips were washed three times with TBST, each wash lasting 10 minutes. Protein bands were then visualized using enhanced chemiluminescence (ECL) substrate (Thermo Scientific) and imaged on an ImageQuant LAS 4000 system. Finally, the intensities of the protein bands were quantified using ImageJ software.

### Protein Enrichment via RNA Pull-Down Assay

RNA-protein interactions involving *circ-MYOCD* were analyzed by RNA pull-down assay. Rat ventricular tissue samples (>500 mg) were fixed with 1% formaldehyde for 10 minutes. The fixation reaction was then quenched by adding 1 M glycine. Following quenching, the tissues were sonicated to generate lysates. Biotinylated probes specifically targeting the *circ-MYOCD* junction site were hybridized with the tissue lysates overnight. Concurrently, streptavidin-coated magnetic beads were pre-washed with hybridization buffer. The pre-washed beads were then incubated with the RNA-biotin probe-lysate mixture for 4 hours at room temperature to capture RNA-bound protein complexes. After incubation, proteins bound to the beads were eluted and subsequently analyzed by mass spectrometry.

### Statistical Analysis

Data are presented as mean ± standard deviation (SD). Comparisons between two groups were performed using unpaired Student’s *t*-test. For experiments involving more than three groups, pairwise comparisons were conducted directly using Tukey’s Honestly Significant Difference (HSD) test. All experiments were repeated with at least three independent biological replicates. Data were visualized using GraphPad Prism software (Version 8.0), and a *p*-value < 0.05 was considered statistically significant for all analyses.

## Results

### SRPK3 expression is upregulated in pathological cardiac hypertrophy

To characterize the expression profile of SRPKs in pathological hypertrophy, we first established an *in vitro* model using neonatal rat cardiomyocytes (NRCMs) treated with phenylephrine (PE; 10, 50, and 100 μM) for 48 hours. Successful induction of hypertrophy was confirmed by the significant upregulation of *ANP* and *Myh7*, alongside the downregulation of *Myh6* mRNA (Figure 1A-B). Subsequent analysis revealed distinct alterations in SRPK isoform expression within hypertrophied NRCMs, with all three SRPKs significantly upregulated following PE treatment (Figure 1C). Tissue profiling across various rat tissues further demonstrated unique expression signatures: while SRPK1 and SRPK2 were ubiquitously expressed, SRPK3 exhibited heart-specific expression (Figure 1D). Notably, within the heart, SRPK3 expression was significantly enriched in primary rat cardiomyocytes compared to cardiac fibroblasts (Figure 1E).

**Figure 1.**
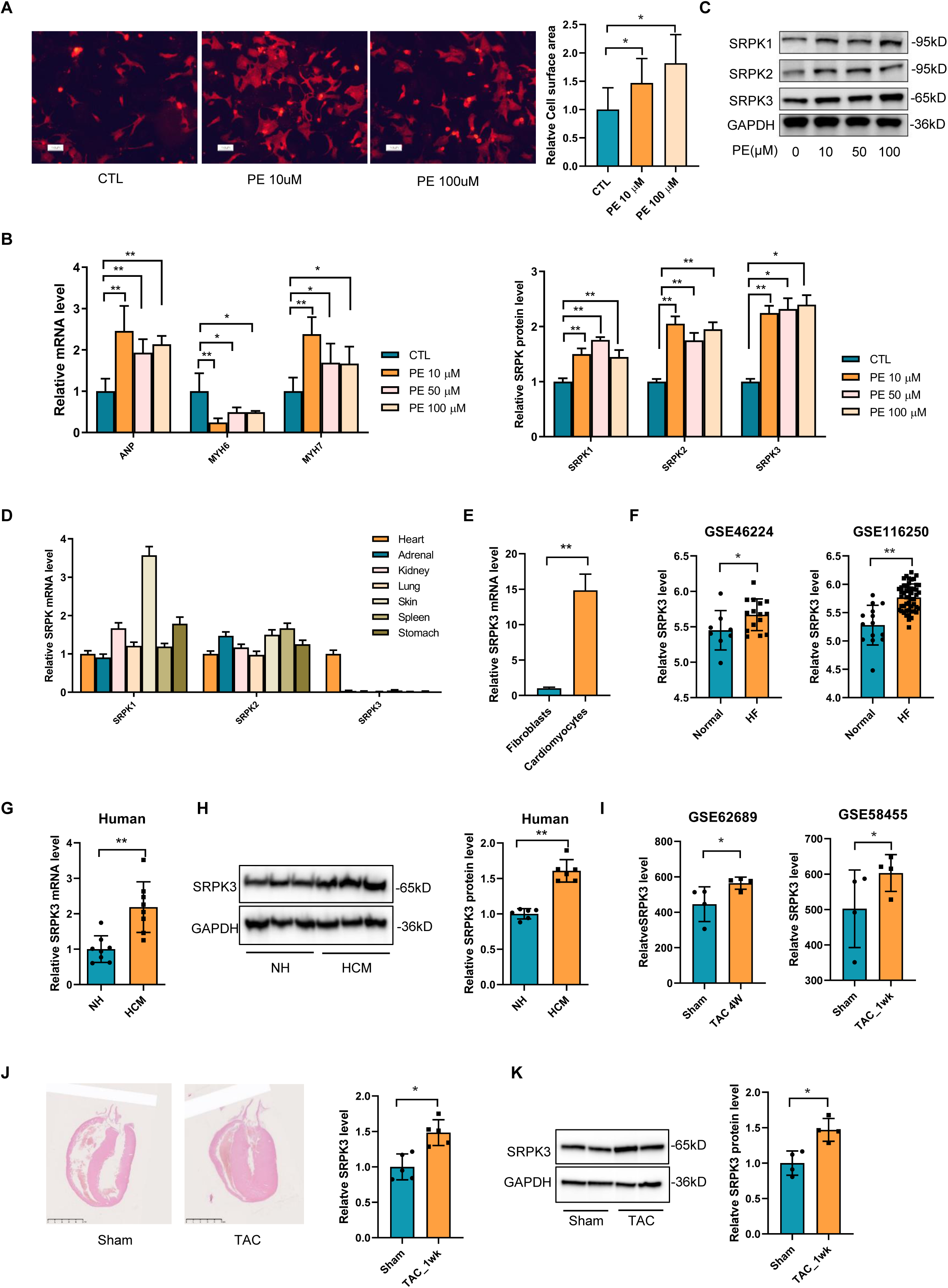
SRPK3 expression is upregulated in pathological cardiac hypertrophy. (A) Induction of hypertrophy in neonatal rat cardiomyocytes (NRCMs) with phenylephrine (10 and 100 μM PE) for 48 h (n=6). (Scale bar: 100μm). (B) Quantitative PCR (qPCR) of hypertrophy markers (*ANP, Myh7*, and *Myh6*) in NRCMs treated PE (n=6). (C) Immunoblotting of SRPK isoforms in NRCMs after hypertrophy induction (n=6). (D) Tissue-specific expression of SRPKs in rat tissues (n=6). (E) SRPK3 expression in primary rat cardiomyocytes (CMs) vs. cardiac fibroblasts (n=4). (F) Meta-analysis of GEO datasets (GSE26887, GSE116250) showing *SRPK3* upregulation in human heart failure versus non-failing controls. (G) qPCR showed that upregulation of *SRPK3* in human HCM hearts (n=8). (H) Immunoblot of SRPK3 protein in human heart tissues (n=6). (I) *SRPK3* mRNA analyzed with GEO dataset about TAC induced cardiac hypertrop hy of mice. (J-K) *SRPK3* mRNA (n=5) and protein levels (n=4) in mice hearts after transverse aortic constriction vs. sham. (J, Scale bar: 5mm). Data: mean ± SD. *\*p* < 0.05, \**\*p* < 0.01.

Translating these findings to human pathology, a meta-analysis of public GEO datasets confirmed robust *SRPK3* upregulation in patients with heart failure (HF) compared to non-failing (NF) controls (Figure 1F). This observation was validated by immunoblotting of human heart tissues, which consistently showed elevated SRPK3 protein levels in hypertrophic samples (Figure 1G-H). Corroborating these clinical and *in vitro* data, *SRPK3* mRNA and protein levels were significantly increased in mouse hearts subjected to transverse aortic constriction (TAC) relative to sham-operated controls (Figure 1I-K).

### SRPK3 drives cardiomyocyte hypertrophy in a kinase-dependent manner

To investigate the functional role of SRPK3 in cardiac hypertrophy, we overexpressed SRPK3 in NRCMs using adenoviral delivery (Ad-SRPK3) (Figure 2A). SRPK3 overexpression resulted in a significant increase in cardiomyocyte surface area, as quantified by α-actinin immunofluorescence staining (Figure 2B), and upregulated the mRNA levels of hypertrophic markers (*ANP*, *BNP*, and *Myh7*) compared to Ad-GFP controls (Figure 2C). Mechanistic interrogation of hypertrophy-related signaling pathways revealed that SRPK3 overexpression specifically activated the SMAD3, FAK, CTNNB, STAT3, and ERK pathways, while exerting no significant effect on mTOR, AKT, P38, or PKA signaling (Figure 2D). Consistent with our *in vitro* findings, *in vivo* adenoviral delivery of SRPK3 (Figure 2E) promoted cardiac hypertrophy (Figure 2G), increase heart weight-to-body weight ratio (HW/BW) (Figure 2F), and elevated expression of *ANP*, *BNP*, and *Myh7* (Figure 2H). Furthermore, bioinformatic analysis of human heart datasets confirmed a positive correlation between SRPK3 and hypertrophic markers (*ANP*, *BNP*, *Myh7*), and a negative correlation with *Myh6* expression (Figure 2I).

**Figure 2.**
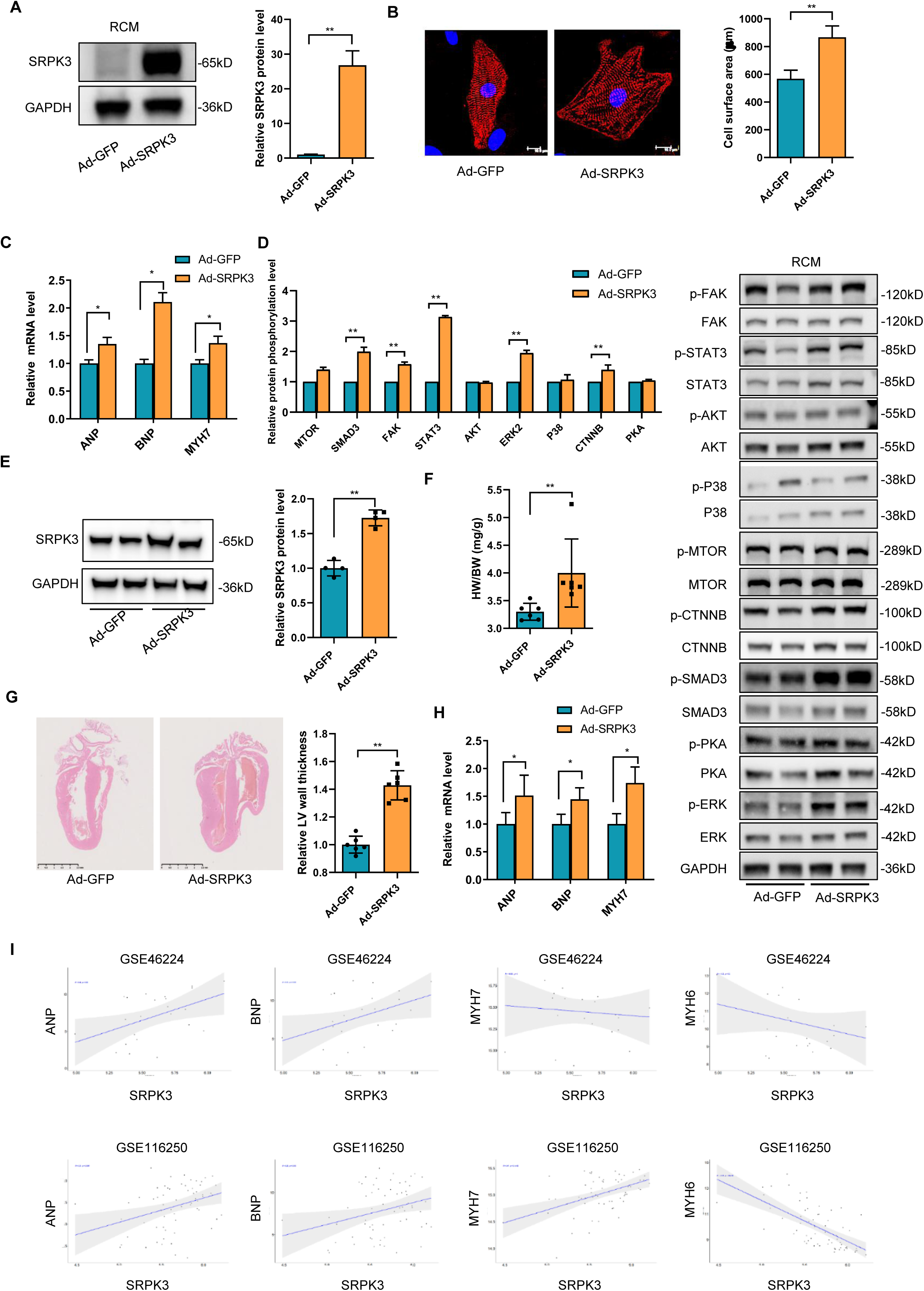
SRPK3 drives cardiomyocyte hypertrophy. (A) Ad-SRPK3 overexpressed SRPK3 protein level in NRCMs (n=4). (B) Representative α-actinin immunofluorescence (scale bar: 10 µm) and cell area quantification (n=10). (C) mRNA levels of *ANP*, *BNP*, and *Myh7* in NRCMs transfected with Ad-SRPK3 in comparison with Ad-GFP (n=6). (D) Transfected with Ad-SRPK3 active cardiac hypertrophic signaling pathways (n=3). (E) Ad-SRPK3 upregulated the protein level of SRPK3 in mice hearts (n=4). (F) Upregulation of SRPK3 protein level increased HW/BW (n=6). (G) Upregulation of SRPK3 protein level promoted ventricular wall thickness by HE staining (n=6) (Scale bar: 5mm). (H) Effects of SRPK3 overexpression on expression of hypertrophy markers in mice hearts (n=6). (I) Correlation of SRPK3 with hypertrophic markers, *ANP*, *BNP*, *Myh7* and *Myh6* in human hearts analyzed with GEO datasets (GSE46224, GSE116250). Data: mean ± SD. *\*p* < 0.05, \**\*p* < 0.01.

To determine whether the pro-hypertrophic function of SRPK3 relies on its enzymatic activity, we analyzed the human SRPK3 protein structure to identify key ATP-binding and activation residues (Figure 3A). We subsequently generated a kinase-dead (KD) SRPK3 mutant adenovirus via targeted mutations (LGWGHFSTV85-93LAWAAFAAA, K108D, and D212A). Immunoblotting confirmed comparable protein expression levels of wild-type (WT) and mutant SRPK3 in transfected NRCMs (Figure 3B). Strikingly, while SRPK3-WT overexpression upregulated hypertrophic markers (*ANP*, *BNP*, *Myh7*) and increased cardiomyocyte size, the kinase-dead mutant failed to elicit these effects (Figure 3C-D). Similarly, SRPK3-WT significantly activated the SMAD3, FAK, CTNNB, STAT3, and ERK signaling pathways, whereas the kinase-dead mutant had no impact on these cascades (Figure 3E). *In vivo* experiments (Figure 3F) further demonstrated that while SRPK3-WT overexpression induced cardiac hypertrophy, as evidenced by hematoxylin-eosin (HE) staining (Figure 3H) and elevated hypertrophic marker expression (Figure 3G), the kinase-dead SRPK3 mutant did not. Collectively, these data establish SRPK3 as a sufficient inducer of cardiomyocyte hypertrophy and demonstrate that its pathological function is strictly kinase-dependent.

**Figure 3.**
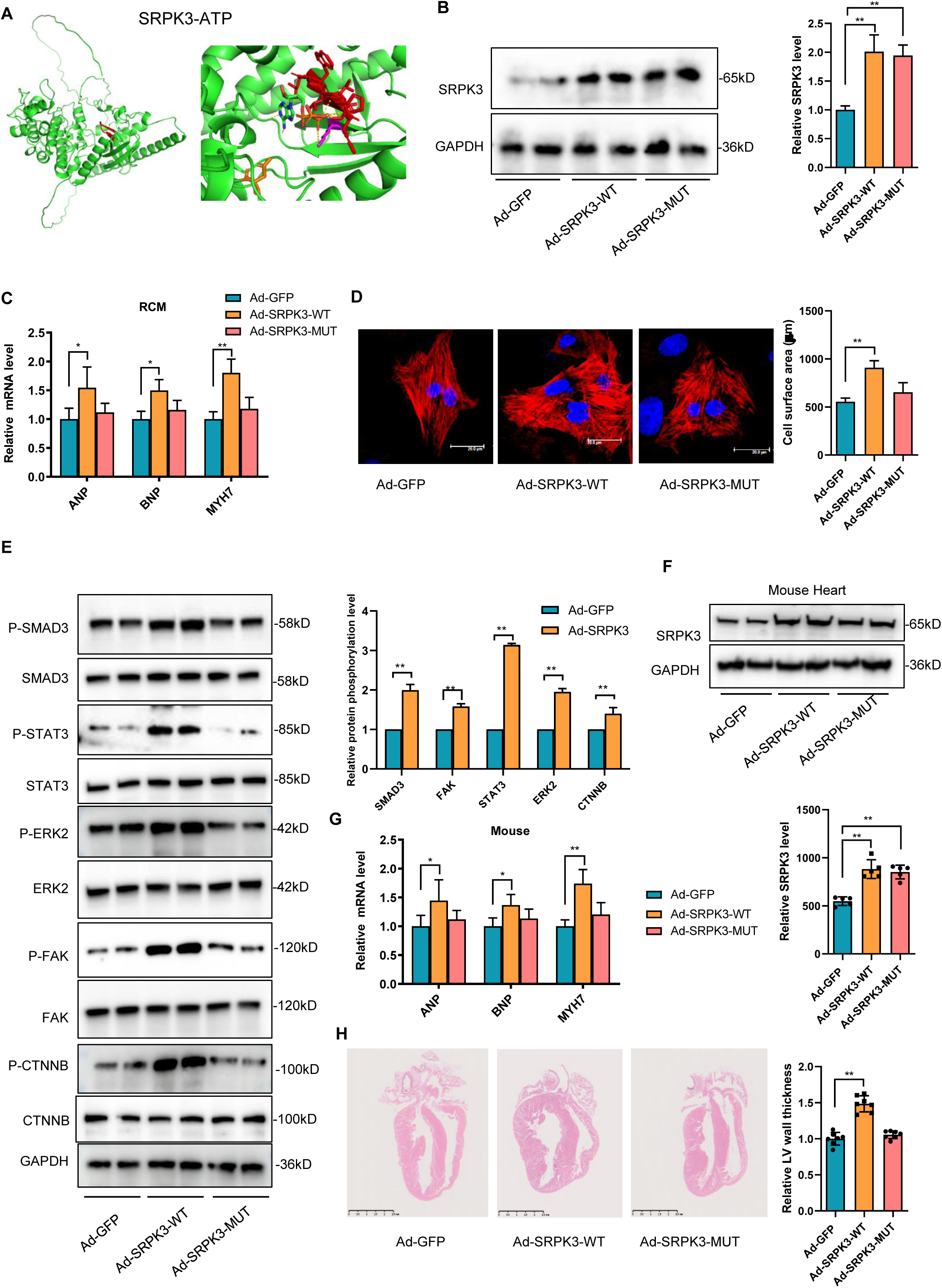
SRPK3 drives cardiomyocyte hypertrophy dependent on its kinase activity. (A) Schematic of SRPK3 wild-type (WT) structure and their binding to ATP substrate. (B) Western blot analyzed the protein level of SRPK3 overexpressed by SRPK3 WT and mutant adenovirus (n=6). (C) qPCR analysis of hypertrophic markers in NRCMs. Ad-SRPK3-WT upregulated, while Kinase Dead mutant failed to upregulated, the expression of *ANP*, *BNP*, and *Myh7* (n=6). (D) Cell surface area in NRCMs transfected with Ad-SRPK3-WT or Ad-SRPK3-KD. Only WT increased cell size (n=8) (Scale bar: 20μm). (E) Transfected with Ad-SRPK3-WT, but not Ad-SRPK3-KD, active cardiac hypertrophic signaling pathways (n=3). (F) *In vivo* adenoviral delivery for SRPK3-WT and SRPK3-KD. WB confirmed the overexpression of SRPK3 in the hearts (n=6). (G) mRNA levels of hypertrophic markers in heart tissues (n=6). (H) H&E staining of heart sections (n=6) (Scale bar: 5mm). Ad-SRPK3-WT induced, while KD did not induced, cardiomyocyte hypertrophy. Data: mean ±SD. *\*p* < 0.05, \**\*p* < 0.01.

### Pharmacological targeting of SRPK3 mitigates hypertrophic remodeling

To evaluate the therapeutic potential of SRPK3, we first employed siRNA-mediated knockdown (Figure 4A). SRPK3 silencing significantly reduced cardiomyocyte size (Figure 4B) and decreased baseline expression of hypertrophy markers (Figure 4C), establishing its necessity for hypertrophic growth. Crucially, SRPK3 knockdown also attenuated phenylephrine (PE)-induced hypertrophy (Figure 4D), as evidenced by the suppression of transcriptional activation of hypertrophy markers (Figure 4E) and the inhibition of cell enlargement (Figure 4F). We further validated these findings pharmacologically using MSC-1186, a pan-SRPK inhibitor (1.2 nM) [21]. Compared to vehicle controls, MSC-1186 treatment significantly reduced cardiomyocyte hypertrophy markers (Figure 4G), and decreased cell surface area (Figure 4H). *In vivo*, MSC-1186 administration effectively suppressed transverse aortic constriction (TAC)-induced cardiac hypertrophy, demonstrated by histological analysis (HE staining, Figure 4J), a reduced heart weight-to-body weight (HW/BW) ratio (Figure 4I), and the normalization of hypertrophic markers (Figure 4K). Collectively, these data demonstrate that MSC-1186 attenuates both baseline and agonist-induced cardiomyocyte hypertrophy.

**Figure 4.**
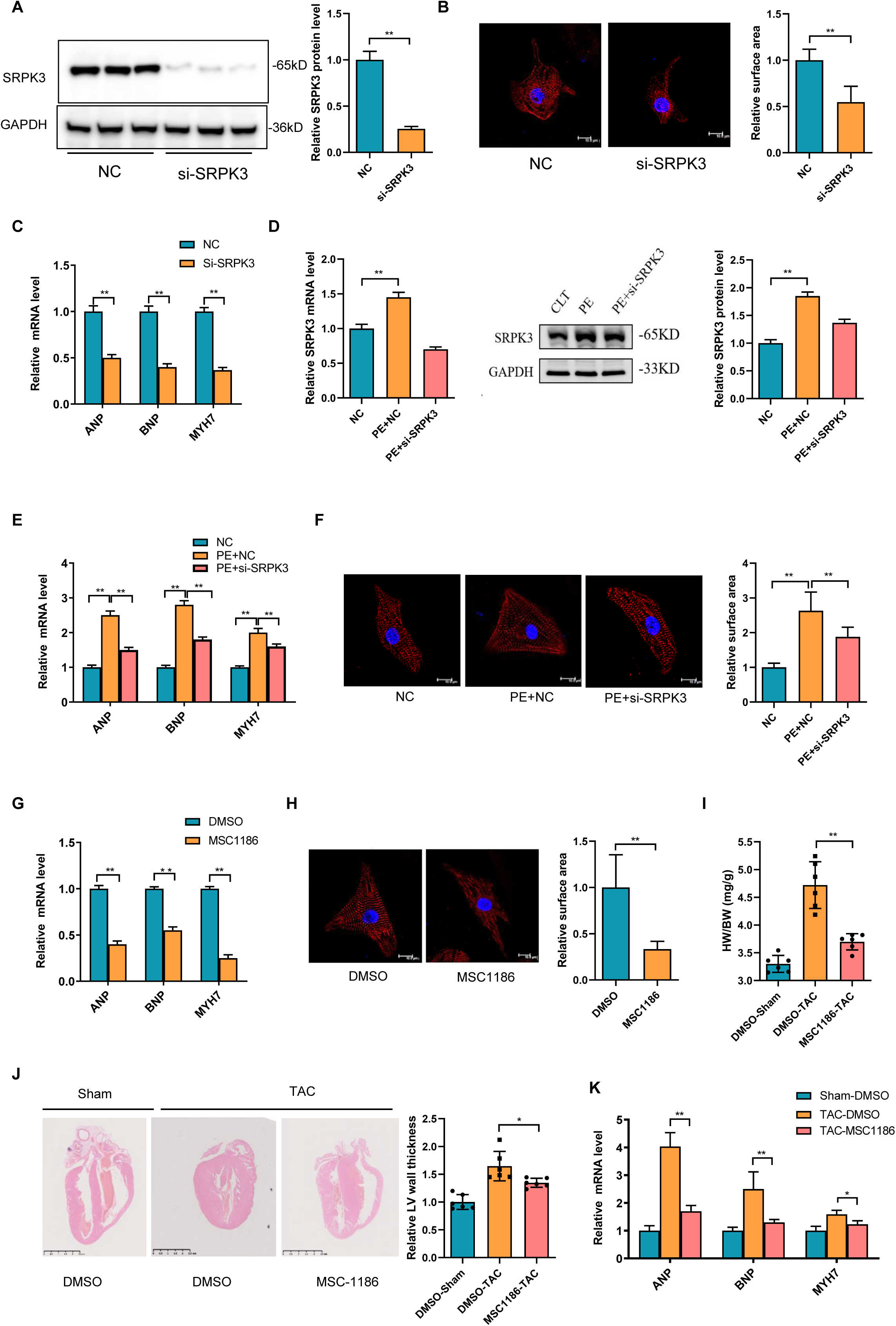
Pharmacological targeting of SRPK3 mitigates hypertrophic remodeling. (A) WB showed siRNA-mediated knockdown of SRPK3 in neonatal rat cardiomyocytes (NRCMs) (n=3). (B) Cell surface area quantification by α-actinin immunofluorescence. SRPK3 knockdown significantly reduced baseline cardiomyocyte size vs. control siRNA (n=10) (Scale bar: 10μm). (C) qPCR analysis showed SRPK3 knockdown downregulated baseline expression of hypertrophy markers (*ANP*, *BNP*, *Myh7*) in NRCMs (n=6). (D-E) SRPK3 siRNA suppressed PE-induced upregulation of hypertrophy markers (*ANP*, *BNP*, *Myh7*) in NRCMs (n=6). (F) Quantification of cell surface area showing SRPK3 knockdown blunted PE-induced cardiomyocyte enlargement (n=6) (Scale bar: 10μm). (G) qPCR analysis of hypertrophy markers in NRCMs treated with pan-SRPK inhibitor MSC-1186 (1.2 nM) (n=6). (H) Cell surface area quantification. MSC-1186 decreased baseline size (n=10) (Scale bar: 10μm). (I) Heart weight-to-body weight ratio (HW/BW) in sham and TAC mice with/without MSC-1186 (n=6). (J) Representative heart sections (H&E staining) from mice subjected to transverse aortic constriction (TAC) and treated with MSC-1186 or vehicle. MSC-1186 suppressed TAC-induced cardiac hypertrophy (n=6) (Scale bar: 5mm). (K) qPCR analysis of cardiac hypertrophic markers (*ANP*, *BNP*, *Myh7*) in heart tissues. MSC-1186 normalized TAC-induced dysregulation (n=6). Data: mean ±SD. *\*p* < 0.05, \**\*p* < 0.01.

### SRPK3 mediates the regulatory back-splicing of *pre-MYOCD* in a kinase-dependent manner

To elucidate the mechanism by which SRPK3 regulates cardiac hypertrophy, we hypothesized that it modulates gene expression via RNA back-splicing. Given the established link between *circRNAs* and heart failure [22, 23], we screened a panel of cardiac-specific *circRNAs*, including *circ-RYR2* (subtypes 14F/9R, 14F/11R, and 35F/34R), *circ-MEF2C*, *circ-CACNA1C*, *circ-MYOCD*, and *circ-SLC8A1*, to determine their sensitivity to SRPK3 levels. We observed that SRPK3 overexpression significantly reduced *circ-MYOCD* levels (Figure 5A) while increasing both *MYOCD* mRNA and protein expression (Figure 5B-C). Conversely, *SRPK3* knockdown upregulated *circ-MYOCD* while downregulating *MYOCD* (Figures 5D-F). In PE-treated cardiomyocytes, *SRPK3* knockdown reversed the PE-induced suppression of *circ-MYOCD* and the concomitant increase in *MYOCD* (Figure 5G-H). Furthermore, pharmacological inhibition with MSC-1186 confirmed that this regulation is kinase activity-dependent: treatment enhanced *circ-MYOCD* expression while suppressing *MYOCD* (Figure 5I-J). These results indicate that SRPK3 modulates *MYOCD* expression by influencing the back-splicing of *MYOCD* precursor RNA.

**Figure 5.**
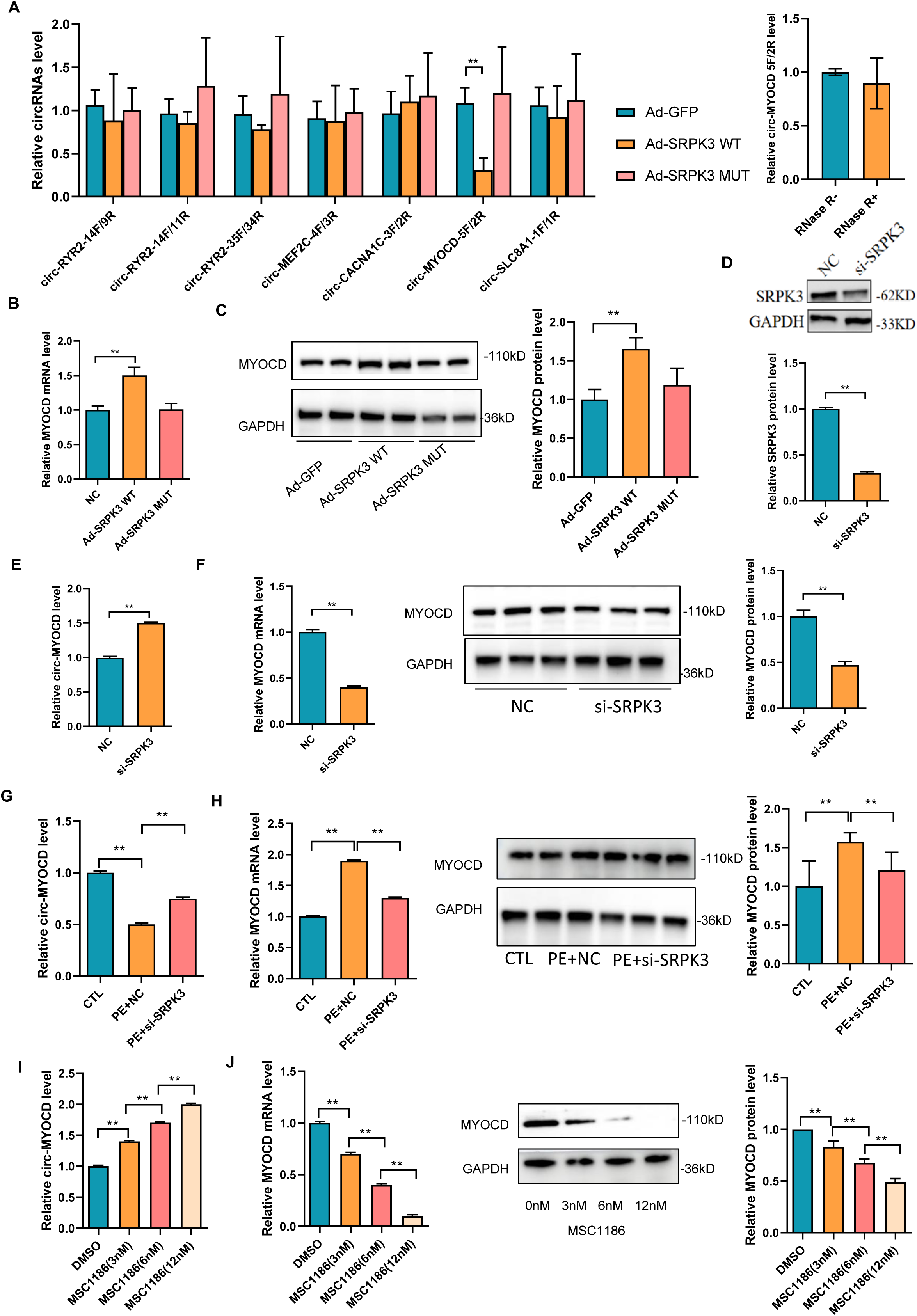
SRPK3 regulates cardiac hypertrophy through kinase-dependent modulation of *circ-MYOCD* back-splicing. (A) Seven cardiac specific *circRNAs* (including *circ-RYR2*, *circ-MEF2C*, *circ-CACNA1C*, *circ-MYOCD*, and *circ-SLC8A1*) were selected to test whether SRPK3 affect backsplicing of their pre-mRNAs. Adenoviral overexpression of SRPK3 (Ad-SRPK3) in cardiomyocytes and its effect on *circ-MYOCD* (subtype 5F/2R). qPCR analysis showed that SRPK3 overexpression significantly reduced *circ-MYOCD* levels (n=4). (B-C) Overexpression of SRPK3 increased *MYOCD* mRNA and protein expression (n=6). (D) SRPK3 knockdown efficiency validated by Western blot (n=6). (E-F) SRPK3 knockdown upregulated *circ-MYOCD* and downregulated *MYOCD* mRNA and protein level (n=6). (G-H) Phenylephrine (PE, 100 μM) suppressed *circ-MYOCD* expression and increase *MYOCD* mRNA and protein in cardiomyocytes, while SRPK3 knockdown reversed this effect (n=6). (I-J) Pharmacological inhibition of SRPK3 with MSC-1186 (1.2 nM) enhanced *circ-MYOCD* and suppressed MYOCD expression (n=6). Data: mean ±SD. *\*p* < 0.05, \**\*p* < 0.01.

### SRPK3 regulates *circ-MYOCD* back-Splicing by directly interacting with and phosphorylating SRSF10

To identify the molecular machinery linking SRPK3 to *circ-MYOCD* regulation, we first performed RNA pull-down assays. Silver staining revealed distinct protein bands enriched in *circ-MYOCD* fractions compared to controls (Figure 6A). Parallel proximity labeling assays (BioID) were conducted to map the SRPK3 interactome. Western blot analysis confirmed the expression of an HA-tagged SRPK3-BirA fusion protein in cardiomyocytes, and co-treatment with biotin yielded streptavidin-detectable biotinylated proteins, validating the labeling of SRPK3-proximal interactors (Figure 6A). Mass spectrometry of the streptavidin pull-down identified a cohort of RNA-binding proteins (RBPs). Intersecting the *circ-MYOCD* interactors with SRPK3-proximal partners highlighted SRSF3, SRSF7, and SRSF10 as candidate regulators (Figure 6B). Functional screening revealed that while SRSF3 and SRSF7 knockdown had no effect (Figures 6D-F), SRSF10 knockdown significantly increased *circ-MYOCD* expression while decreasing *Myocd* mRNA (Figures 6G-H). Conversely, SRSF10 overexpression suppressed *circ-MYOCD* but increased *Myocd* mRNA (Figures 6I–J). Mechanistically, SRSF10 RIP assays confirmed a direct interaction with *circ-MYOCD* (Figure 6K). Sequence analysis identified three potential SRSF10 binding motifs (Figure 6L), and motif-specific RIP assays pinpointed the F3R3 region as the functional binding site (Figure 6M).

**Figure 6.**
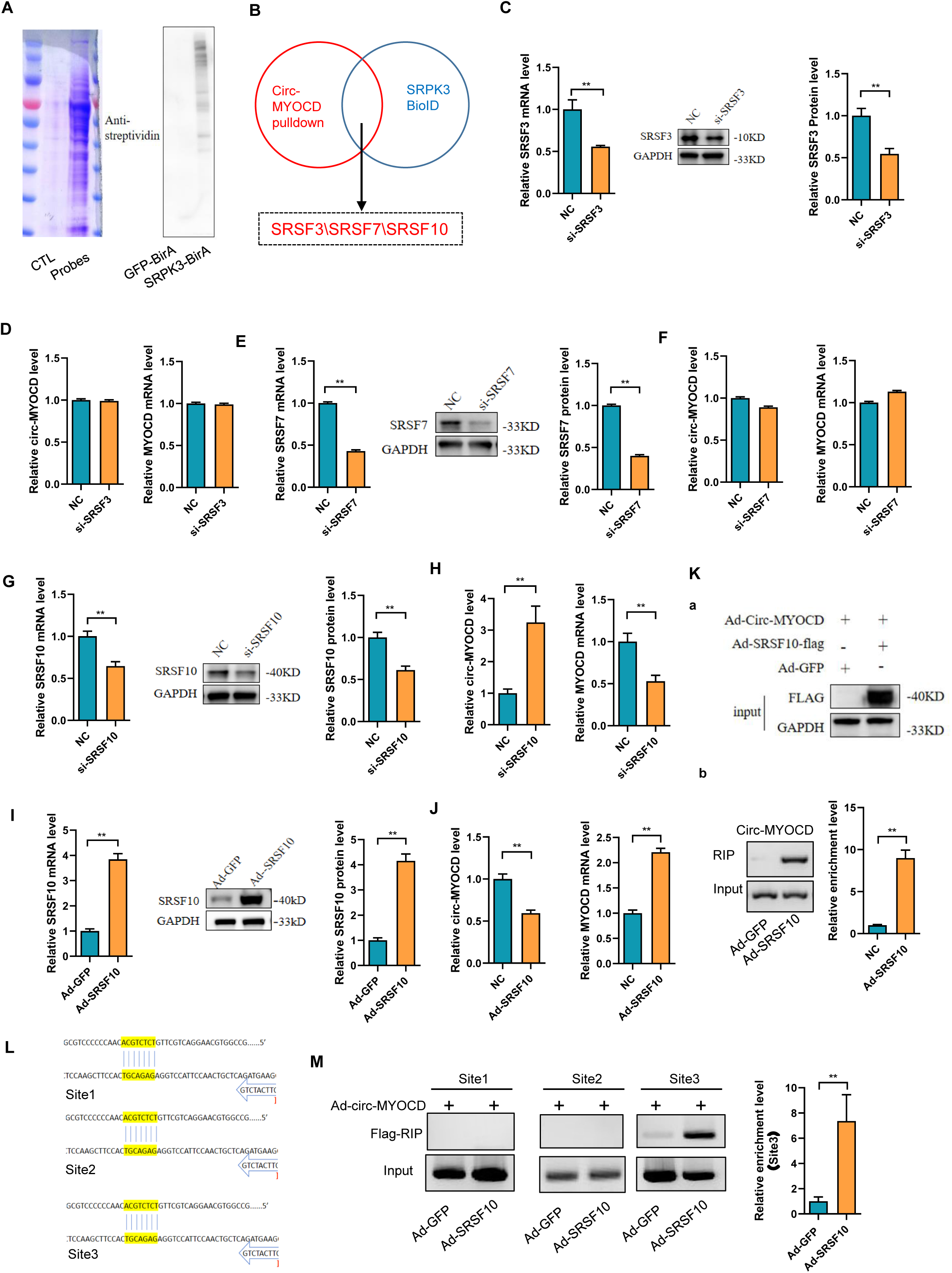
SRSF10 mediates SRPK3-dependent regulation of *circ-MYOCD* biogenesis. (A-B) RNA pull-down assay for *circ-MYOCD*-interacting proteins. Coomassie staining of SDS-PAGE-separated proteins from neonatal rat cardiomyocytes (NRCMs) shows distinct bands in *circ-MYOCD*-enriched fractions (lane 2) vs. control (lane 1). Validation of SRPK3 proximity labeling. Western blot of HA-tagged SRPK3-BirA fusion protein in Ad-SRPK3-BirA-HA-transfected NRCMs. Biotin treatment induced streptavidin-detectable biotinylation of SRPK3-proximal interactors. (B) Venn diagram of proteins co-precipitated with *circ-MYOCD* (RNA pull-down) and SRPK3 (BioID). SRSF3, SRSF7, and SRSF10 are shared RNA-binding proteins (RBPs). (C-F) qPCR analysis of *circ-MYOCD* and *MYOCD* mRNA in NRCMs after siRNA-mediated knockdown of SRSF3 or SRSF7. No significant changes were observed (n=6). (G-H) SRSF10 knockdown increased *circ-MYOCD* and decreased *MYOCD* mRNA (n=6). (I-J) SRSF10 overexpression decreased *circ-MYOCD* and increased *MYOCD* mRNA (n=6). (K) RNA immunoprecipitation (RIP)-qPCR confirms direct binding between SRSF10 and *circ-MYOCD* (n=6). (L) *circ-MYOCD* fragments contains 3 candidate SRSF10 binding motifs and design primer pairs to detect them (F1R1, F2R2, F3R3). (M) RIP-qPCR of SRSF10 specifically enriched the F3R3 motif region (n=6). Data: mean ±SD. *\*p* < 0.05, \**\*p* < 0.01.

We next investigated how SRPK3 regulates SRSF10. BioID proximity labeling followed by Western blot showed that SRPK3-BirA biotinylates SRSF10 (Figure 7A), indicating close *in vivo* proximity. This physical interaction was corroborated by reciprocal co-immunoprecipitation (Co-IP) assays (Figure 7B). Furthermore, SRPK3 overexpression significantly increased global protein phosphorylation (Figure 7C) and specifically enhanced SRSF10 phosphorylation (Figure 7D). Structural analysis predicted serine residues 129, 131, and 133 (S129/131/133) as potential phosphorylation sites (Figure 7E). To test their functional significance, we generated phospho-deficient SRSF10 mutants (S129/131/133A). Strikingly, these mutations abolished the ability of SRSF10 to suppress *circ-MYOCD* biogenesis and promote linear *MYOCD* production (Figure 7F-G). Consistently, while wild-type SRSF10 increased MYOCD protein levels, the phospho-deficient mutant failed to do so. Thus, SRPK3-mediated phosphorylation at these specific serines is required for SRSF10’s suppressive effect on *circ-MYOCD* formation.

**Figure 7.**
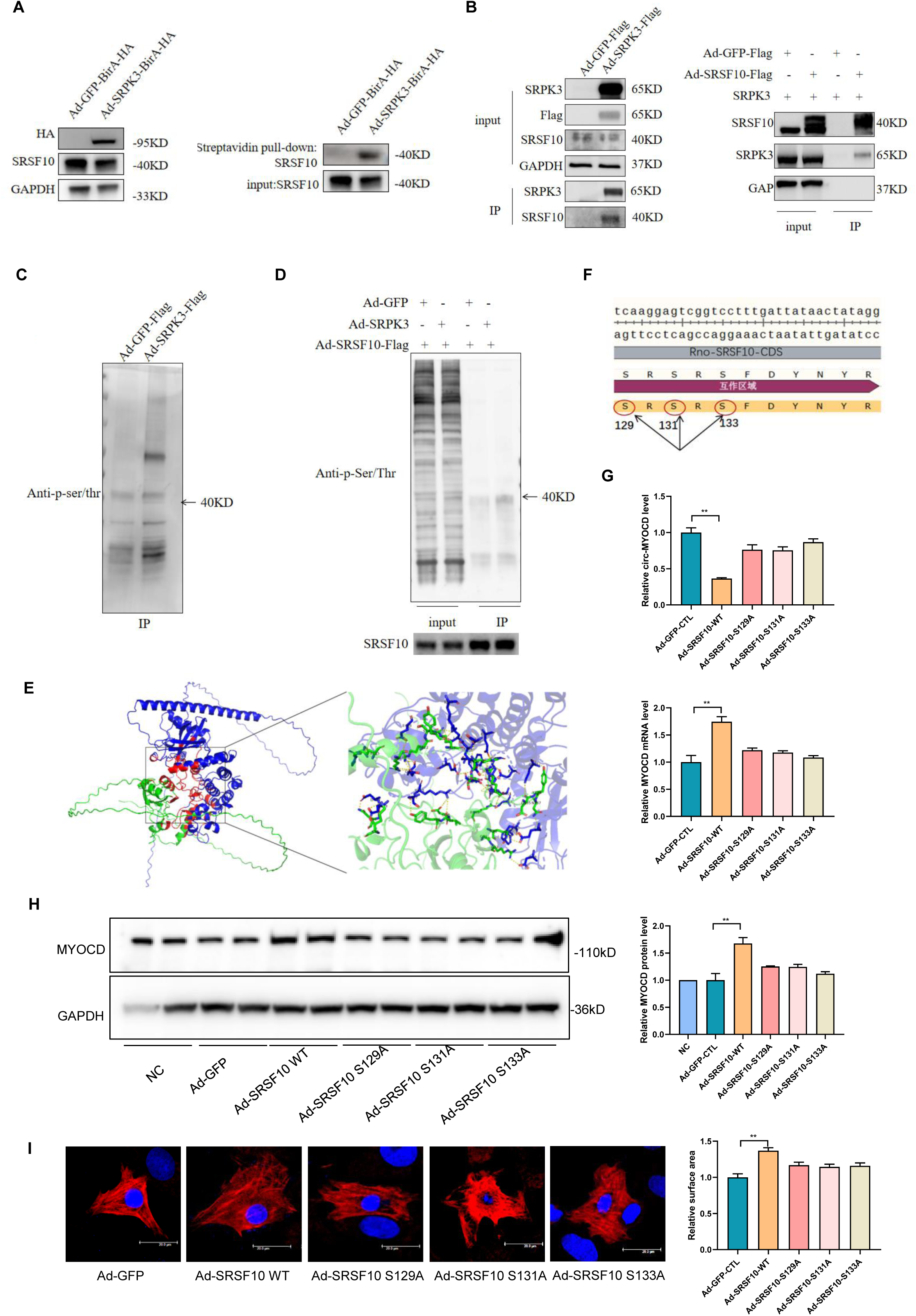
SRPK3 phosphorylates SRSF10 to suppress *circ-MYOCD* biogenesis. (A) Proximity labeling validation. Western blot of biotinylated SRSF10 in SRPK3-BirA-expressing NRCMs with biotin treatment. (B) Co-immunoprecipitation (Co-IP) of SRPK3 and SRSF10. Endogenous SRSF10 co-precipitated with Flag-tagged SRPK3 (top), and SRPK3 co-precipitated with SRSF10 antibodies. (C) Global protein phosphorylation in SRPK3-overexpressing NRCMs. (D) SRSF10 phosphorylation assay. SRPK3 overexpression enhanced SRSF10 phosphorylation. (E) Predicted SRPK3-SRSF10 interaction structure. Serine residues S129, S131, S133 (red) are potential phosphorylation sites. (F-H) Functional impact of SRSF10 phosphorylation mutants. Adenoviral expression of phospho-deficient SRSF10 (S129/131/133A) in NRCMs failed to suppress *circ-MYOCD* or downregulate *MYOCD* mRNA and protein, unlike wild-type SRSF10 (n=3). (I) Adenoviral expression of phospho-deficient SRSF10 (S129/131/133A) in NRCMs failed to induced cardiomyocyte hypertrophy (n=10) (Scale bar: 20μm). Data: mean ±SD. *\*p* < 0.05, \**\*p* < 0.01.

### Downregulation of *Circ-MYOCD* leads to cardiac hypertrophy

RT-PCR showed that *circ-MYOCD* was specifically expressed in rat heart (Figure 8A). In a rat model of myocardial hypertrophy induced by PE and Ang II (Figure 8B), expression analysis revealed significant downregulation of *circ-MYOCD* (5F/2R). Further validation in the transverse aortic constriction (TAC) model (Figure 8C) and human failing hearts (Figure 8D) confirmed significant downregulation of *circ-MYOCD* compared to controls.

**Figure 8.**
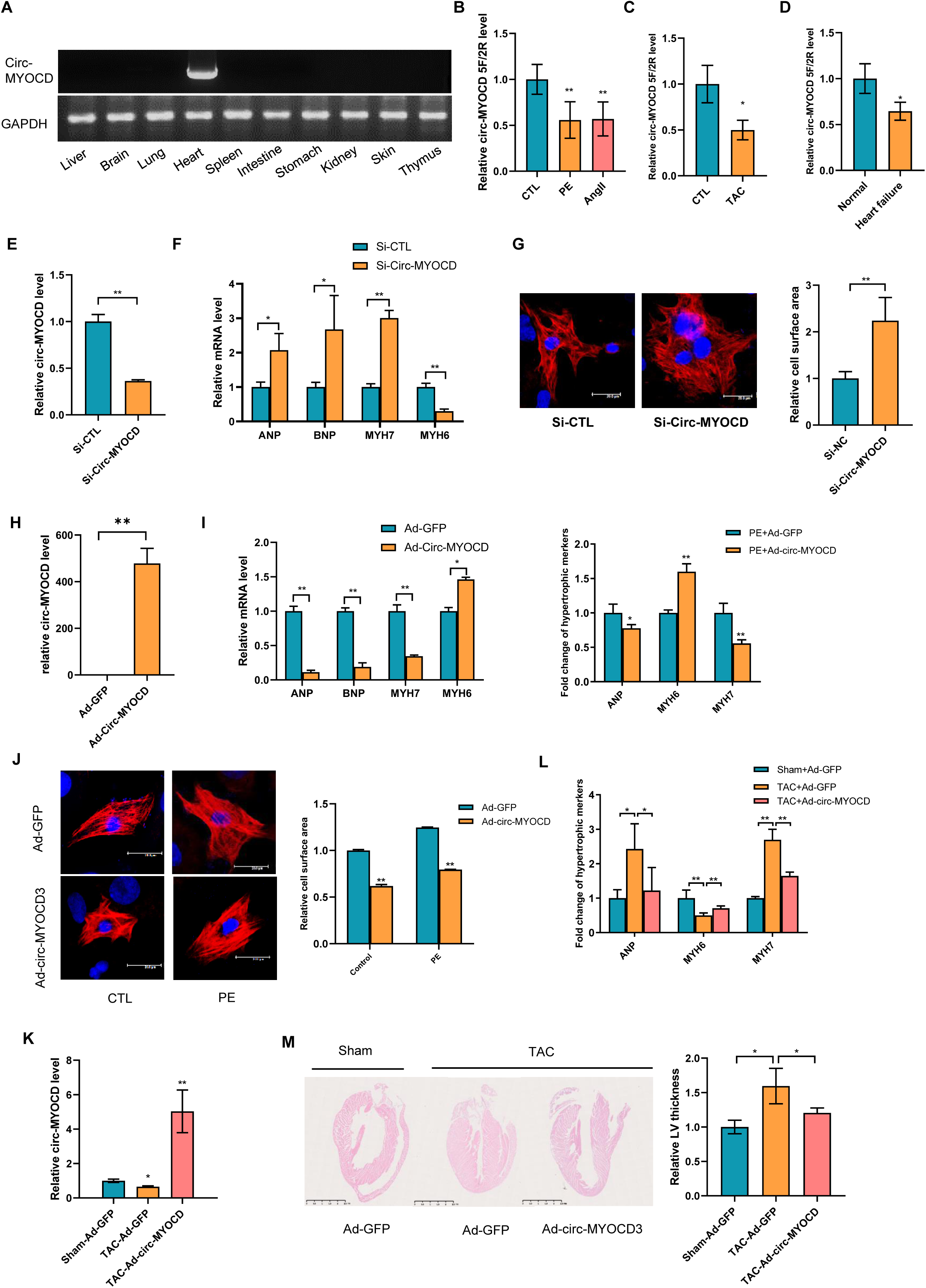
*Circ-MYOCD* protects against cardiac hypertrophy. (A) Tissue-restricted expression of *circ-MYOCD* in rat organs by RT-PCR, showing exclusive myocardial enrichment. (B) Significant downregulation of *circ-MYOCD* (5F/2R) in PE-treated cardiomyocytes versus controls. (C) Significant *circ-MYOCD* downregulation in pressure-overloaded hearts versus sham controls. (D) Conserved downregulation of *circ-MYOCD* in human failing hearts (HF) versus non-failing (NF) controls. (E) siRNA-mediated knockdown of *circ-MYOCD* in neonatal rat cardiomyocytes (NRCMs) (n=6). (F) qPCR analysis of hypertrophy markers (*ANP*, *BNP*, *Myh7*) and *Myh6*. *circ-MYOCD* knockdown upregulated *ANP*, *BNP*, and *Myh7* while downregulating *Myh6* (n=6). (G) Cardiomyocyte surface area quantified by confocal microscopy of cardiac troponin T (CTNT)-stained cells. *circ-MYOCD* knockdown increased basal cell size (n=10) (Scale bar: 20μm). (H-I) qPCR analysis of hypertrophy markers in NRCMs overexpressing *circ-MYOCD* via adenovirus (Ad-*circ-MYOCD*). *circ-MYOCD* overexpression decreased *ANP*, *BNP*, and *Myh7* while restoring *Myh6* (n=6) under basal and PE-stimulated conditions. (J) Cell surface area measurement. *circ-MYOCD* overexpression reduced cardiomyocyte size (n=10) (Scale bar: 20μm). (K) Intracardiac Ad-*circ-MYOCD* injection followed by transverse aortic constriction (TAC). (L) *Circ-MYOCD* overexpression attenuated TAC-induced *ANP*/*Myh7* upregulation and *Myh6* downregulation (M) HE staining demonstrated reduced left ventricular wall thickening in TAC+Ad-*circ-MYOCD* hearts. Data: mean ± SEM; n=3-5 (cells), n=6-8 (mice). \**p* < 0.05, \*\**p* < 0.01.

To define the functional role of *circ-MYOCD*, we silenced it in cardiomyocytes using specific siRNAs (Figure 8E). *Circ-MYOCD* knockdown significantly upregulated hypertrophy markers (*ANP*, *BNP*, and *Myh7*), downregulated *Myh6* (Figure 8F), and markedly increased cardiomyocyte surface area (Figure 8G), suggesting a cardioprotective role. Conversely, *circ-MYOCD* overexpression (Ad-*circ-MYOCD*) suppressed *ANP*, *BNP*, and *Myh7* expression while restoring *Myh6* levels (Figure 8H-I) and significantly reducing cell surface area compared to GFP controls even under PE stimulation (Figure 8J).

Following intracardiac injection of a circ-MYOCD-overexpressing adenovirus in mice, transverse aortic constriction (TAC) surgery was performed to establish a cardiac hypertrophy model. One week after TAC surgery, quantitative PCR analysis of cardiac tissue revealed that *circ-MYOCD* overexpression significantly attenuated the TAC-induced upregulation of hypertrophy markers *ANP* and *Myh7*, while significantly reversing the TAC-induced downregulation of *Myh6* expression (Figure 8K-L). Furthermore, histological assessment by HE staining demonstrated that *circ-MYOCD* overexpression significantly reduced TAC-induced left ventricular wall thickening (Figure 8M).

### *Circ-MYOCD* Regulates Chaperonin and Heat Shock Protein Networks Essential for Proteostasis

Mechanistically, proteins interacting with *circ-MYOCD* were identified using RNA pull-down assays. Coomassie Brilliant Blue staining of SDS-PAGE-separated proteins revealed distinct bands specifically enriched in the *circ-MYOCD* pull-down fraction compared to controls (Figure 9A). Mass spectrometry analysis confirmed the enrichment of cytoplasmic proteins, including molecular chaperones and heat shock proteins (HSPs), suggesting their functional relevance. Protein-protein interaction network analysis using STRING highlighted specific interactions among key players like HSPA5, HSPA9, HSP90AA1, HSP90B1, CCT6A,and CCT4 (Figure 9B). These results was consistent with the predominant cytoplasmic localization of *circ-MYOCD* (Figure 9C).

**Figure 9.**
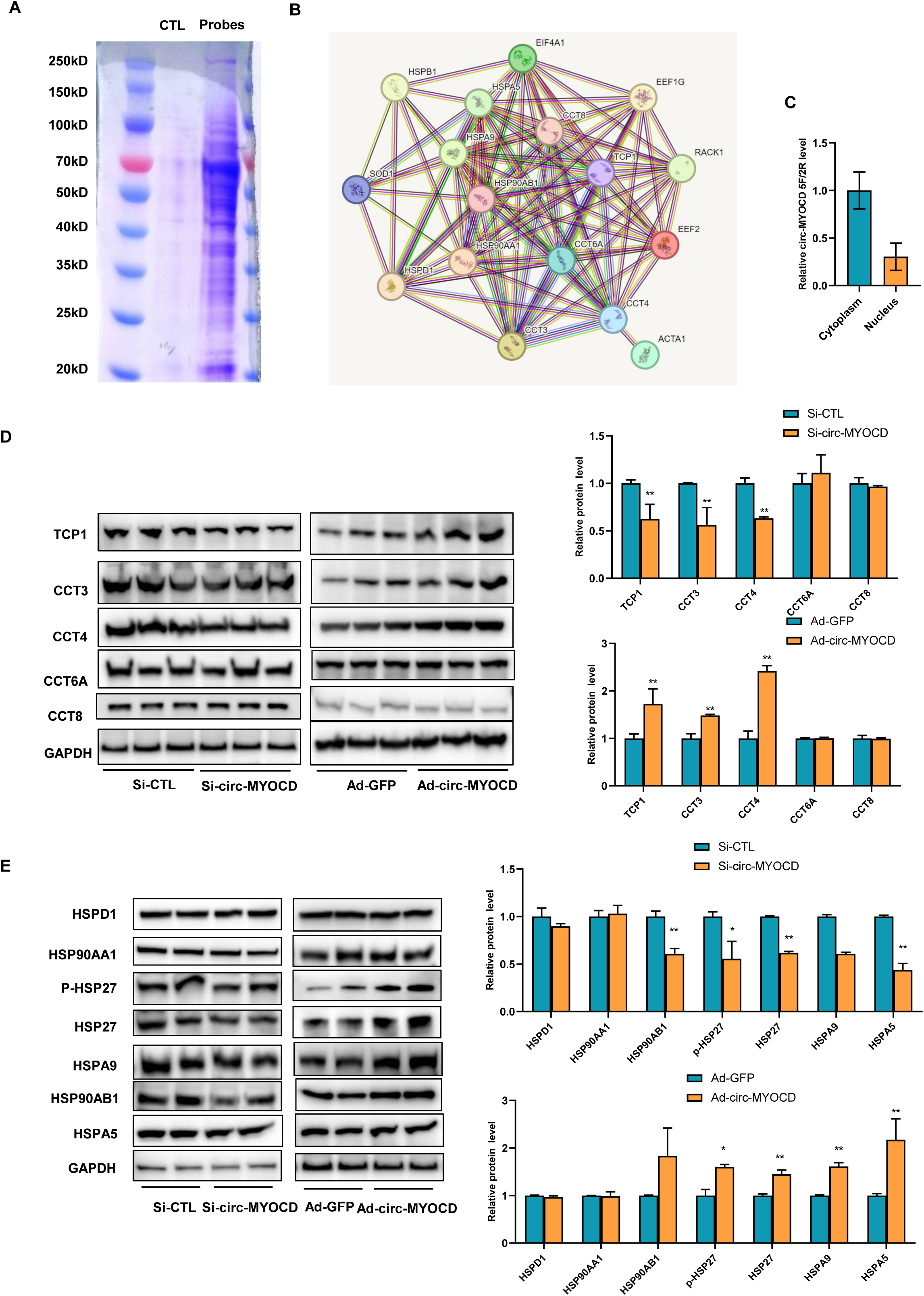
*Circ-MYOCD* orchestrates proteostasis by regulating TRiC complex components and heat shock proteins. (A) RNA pull-down assay identifies *circ-MYOCD*-binding proteins. Silver staining of SDS-PAGE-separated proteins reveals distinct bands enriched in *circ-MYOCD* pull-down fractions versus negative oligos. (B) Protein interaction network of core *circ-MYOCD* partners. STRING database analysis highlights physical/functional interactions among enriched chaperones including HSPA9, HSP90AA1, HSP90B1, and CCT. (C) Nucleoplasmic fractionation of rat cardiomyocytes followed by qRT-PCR reveals predominant cytoplasmic localization of *circ-MYOCD*. (D) *Circ-MYOCD* knockdown significantly downregulated TCP1, CCT3, and CCT4 protein levels, while overexpression selectively upregulated these TRiC components. (E) *Circ-MYOCD* depletion reduced HSPA9, GRP78, and HSP90AB1 expression, with reciprocal upregulation upon overexpression. Notably, phosphorylated HSP27 (p-HSP27) levels decreased with knockdown and increased with overexpression, while total HSP27 remained unchanged. \**p* < 0.05, \*\**p* < 0.01 vs. respective controls.

Knockdown of *circ-MYOCD* significantly downregulated multiple components of the chaperonin TCP1 ring complex (TRiC), including TCP1, CCT3, and CCT4. Conversely, *circ-MYOCD* overexpression selectively upregulated TCP1, CCT3, and CCT4 (Figure 9D). Similarly, *circ-MYOCD* depletion reduced the levels of HSPA9, GRP78, and HSP90AB1, while its overexpression increased the levels of these HSPs. Notably, phosphorylated HSP27 (p-HSP27) levels decreased upon *circ-MYOCD* knockdown and increased upon overexpression, while total HSP27 protein levels remained unchanged (Figure 9E). These results establish *circ-MYOCD* as a master regulator of proteostatic machinery. It differentially orchestrates the expression of specific TRiC subunits and HSP isoforms and uniquely promotes HSP27 phosphorylation. This implicates *circ-MYOCD* as a potential therapeutic target for modulating cardiac stress responses.

### *Circ-MYOCD* counteracts SRPK3 function to protect against cardiac hypertrophy

Knockdown of *circ-MYOCD* specifically activated SMAD3, STAT3, and ERK signaling pathways, without affecting mTOR, AKT, P38, PKA, FAK, or CTNNB signaling (Figure 10A). Overexpression of *circ-MYOCD* did not significantly affect the phosphorylation of mTOR, SMAD3, AKT, ERK2, P38, CTNNB1, or PKCα. These complementary loss- and gain-of-function experiments demonstrate that *circ-MYOCD* acts as a selective modulator of hypertrophic signaling pathways. Upon its loss, *circ-MYOCD* potentiates SMAD3, STAT3, and ERK2 signaling activity while inhibiting FAK phosphorylation. Upon its overexpression, it enhances FAK phosphorylation while suppressing STAT3 phosphorylation. Critically, *circ-MYOCD* does not alter other major pathways (PI3K/AKT, mTOR, P38 MAPK, Wnt/β-catenin, PKC), highlighting its specific role in regulating the FAK and JAK/STAT3 nodes, alongside SMAD2 and ERK2 activity (Figure 10B).

**Figure 10.**
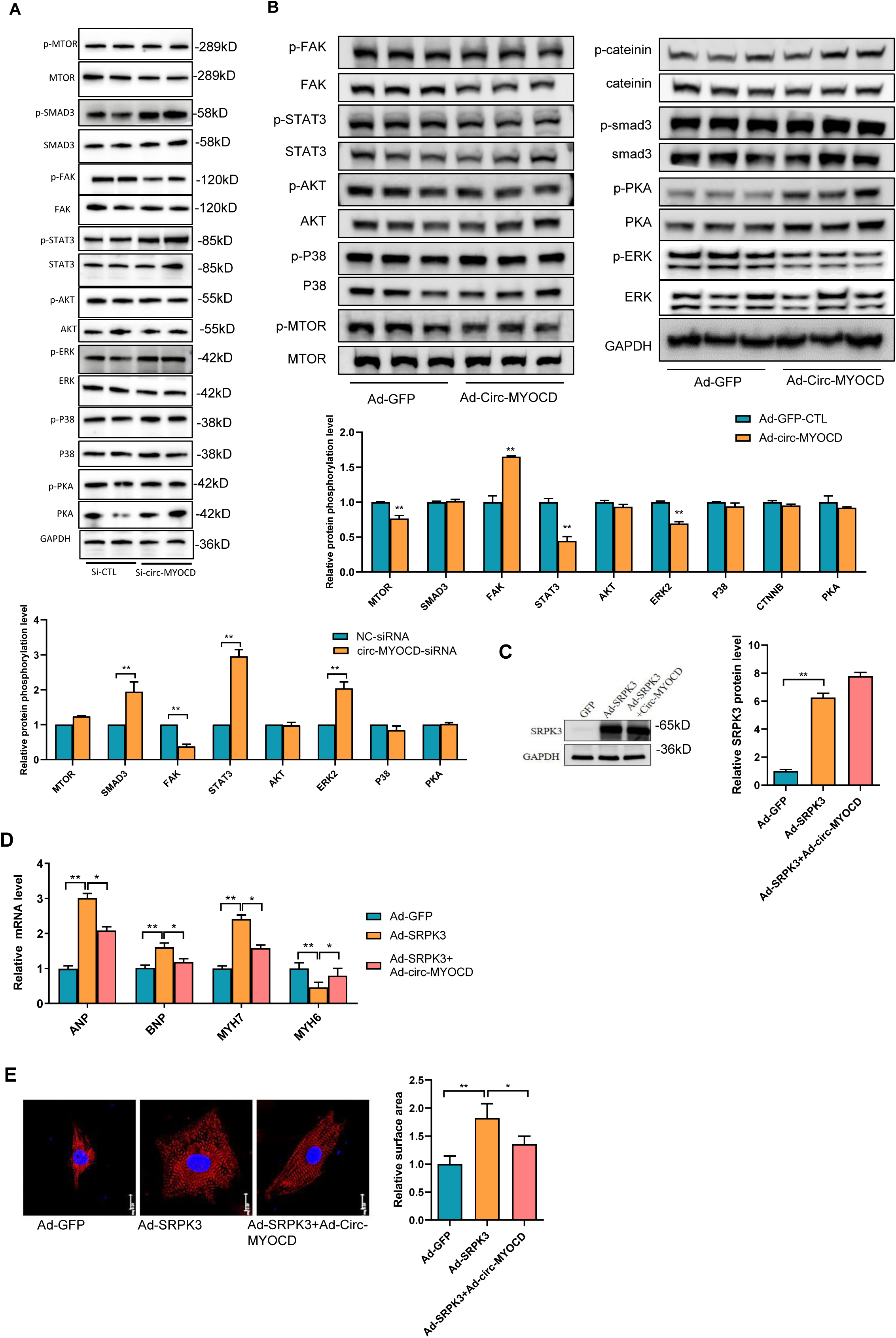
*circ-MYOCD* antagonizes SRPK3-driven cardiac hypertrophy by modulating specific signaling pathways. (A) Loss-of-function analysis in neonatal rat cardiomyocytes revealed that *circ-MYOCD* knockdown significantly increased phosphorylation of SMAD3 (TGF-β pathway), STAT3 (JAK/STAT pathway), and ERK2 (MAPK/ERK pathway) while reducing FAK phosphorylation compared to non-targeting control siRNA (NC-siRNA). Phosphorylation of mTOR, AKT (PI3K/AKT pathway), P38 (MAPK pathway), CTNNB (Wnt/β-catenin pathway), and PKA (cAMP pathway) remained unchanged. (B) *Circ-MYOCD* overexpression significantly enhanced FAK phosphorylation and suppressed STAT3 phosphorylation, without altering mTOR, SMAD3, AKT, ERK2, P38, CTNNB1, or PKCα phosphorylation. (C-D) qPCR analysis of hypertrophy markers in SRPK3-overexpressing NRCMs with/without *circ-MYOCD* restoration. *circ-MYOCD* rescued SRPK3-induced upregulation of *ANP*, *BNP*, and *Myh7* (n=6). (E) Cell surface area quantification. *circ-MYOCD* attenuated SRPK3-induced cardiomyocyte enlargement (n=10) (Scale bar: 10μm). Data: mean ±SD. *\*p* < 0.05, **p < 0.01.

Finally, to determine if *circ-MYOCD* antagonizes SRPK3-driven hypertrophy, we restored its expression in SRPK3-overexpressing cardiomyocytes (Figure 10C). While SRPK3 overexpression upregulated hypertrophy markers (Figure 10D) and increased cell size (Figures 10E), the restoration of *circ-MYOCD* effectively rescued this pathological remodeling (Figures 10C-E).

### Hierarchical regulation of pathological cardiac remodeling through the SRPK3/*circ-MYOCD*/MYOCD axis

Collectively, our findings delineate a cardiac-specific regulatory axis wherein SRPK3 governs pathological remodeling by suppressing the cardioprotective circular RNA, *circ-MYOCD*. This framework operates through three interconnected mechanistic tiers. First, the pathological upregulation of SRPK3 observed in human heart failure and experimental hypertrophy models drives the phosphorylation of the splicing factor SRSF10. Second, this phosphorylation event inhibits the back-splicing of exon 2-5, a process essential for *circ-MYOCD* biogenesis. This kinase-dependent suppression initiates a feed-forward cascade: the resulting reduction in *circ-MYOCD* relieves its competitive inhibition on the linear splicing of *MYOCD* pre-mRNA, thereby promoting the production of linear *MYOCD* mRNA. Consequently, MYOCD accumulation drives the transcriptional activation of the ERK, STAT3, and SMAD3 signaling pathways, ultimately inducing pathological hypertrophy (Figure 11).

**Figure 11.**
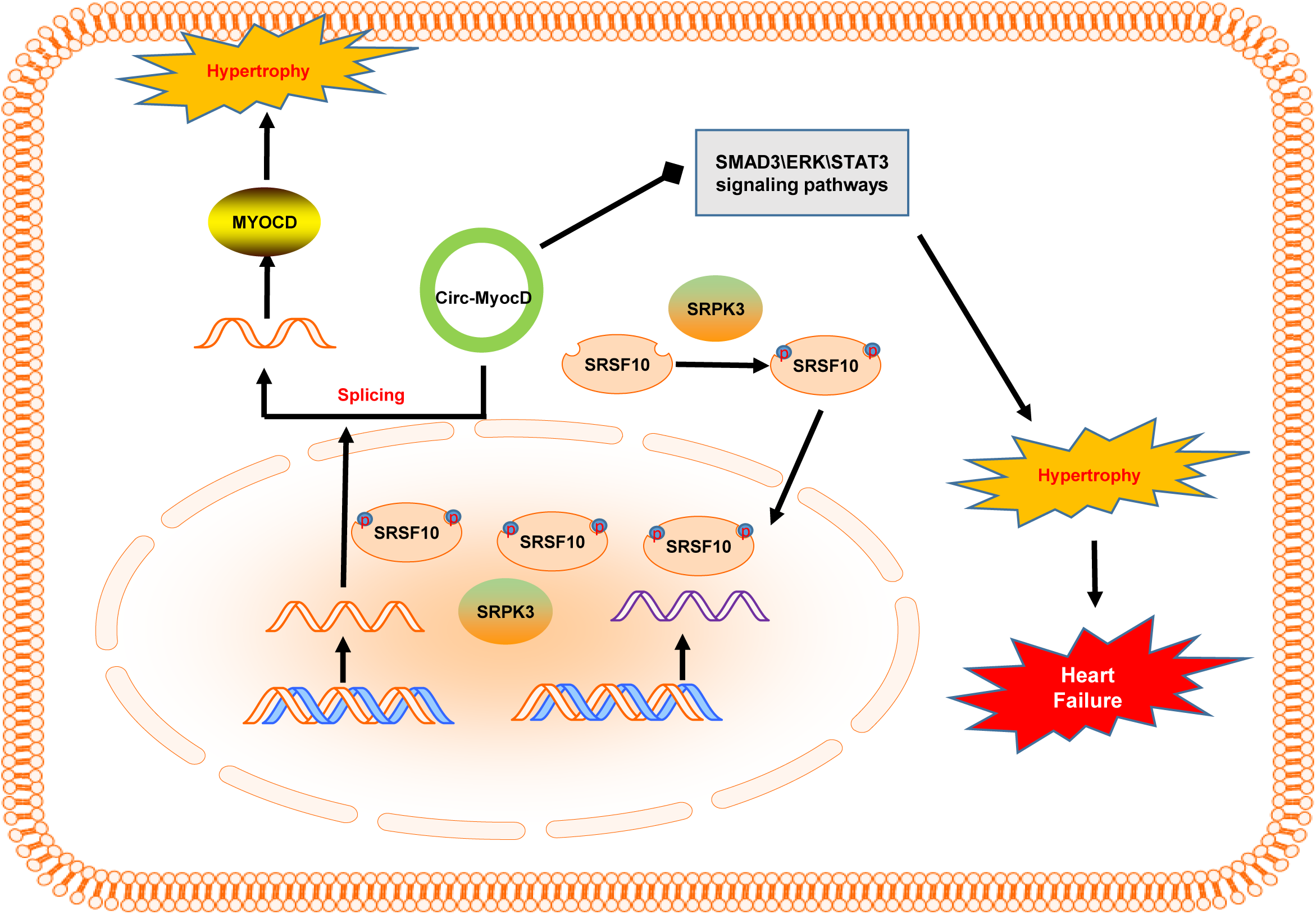
Hierarchical regulation of pathological cardiac remodeling through the SRPK3/*circ-MYOCD*/MYOCD axis. Pathological upregulation of SRPK3 in heart failure phosphorylates the splicing factor SRSF10, which suppresses exon 2-5 back-splicing necessary for *circ-MYOCD* and MYOCD biogenesis. Reduced *circ-MYOCD* levels relieve its competitive inhibition on linear splicing of *MYOCD* pre-mRNA, resulting in increased MYOCD protein production. MYOCD drives transcriptional activation of pro-hypertrophic signaling, while concurrent *circ-MYOCD* depletion disrupts its scaffolding function. SRPK3 elevation in human heart failure and experimental models promotes *circ-MYOCD* suppression, which amplifies MYOCD/ERK activation and proteostasis collapse, collectively accelerating pathological hypertrophy. Arrows denote sequential causality.

## Discussion

Cardiac hypertrophy represents a maladaptive response to sustained hemodynamic stress, frequently progressing to heart failure, a leading cause of global mortality [3]. While transcriptional regulators like ERK and STAT3 are established drivers of this pathology, the contribution of post-transcriptional mechanisms, particularly RNA splicing and circRNA biogenesis, has remained underexplored. Our study addresses this gap by identifying SRPK3, a striated muscle-specific kinase, as a master regulator of pathological cardiac remodeling.

Unlike the ubiquitously expressed SRPK1 and SRPK2, SRPK3 is exclusively enriched in striated muscle, aligning with its specialized role in cardiac stress adaptation. We observed consistent SRPK3 upregulation across human heart failure transcriptomes, *in vivo* transverse aortic constriction (TAC) models, and *in vitro* phenylephrine (PE)-stimulated cardiomyocytes. Functional studies confirmed its causal role: SRPK3 overexpression amplified hypertrophic markers (*ANP*, *BNP*, *Myh7*) and cardiomyocyte enlargement, whereas siRNA-mediated knockdown or pharmacological inhibition with MSC-1186 significantly attenuated remodeling. This distinct tissue specificity positions SRPK3 as a highly promising therapeutic target with minimized off-target potential.

Mechanistically, we elucidate a precise hierarchical pathway where SRPK3 phosphorylates the splicing factor SRSF10. This phosphorylation event is critical, as it leads to the suppression of *circ-MYOCD* biogenesis while enhancing the efficiency of canonical splicing. The reduction in *circ-MYOCD* levels results in a concomitant increase in linear *MYOCD* expression. This shift in the linear-to-circular ratio is functionally significant; the depletion of *circ-MYOCD* liberates ERK, STAT3, and SMAD3 signaling, thereby accelerating hypertrophy. Conversely, *circ-MYOCD* overexpression effectively reverses SRPK3-driven pathology, confirming its role as a negative feedback regulator.

A critical insight from our study is the competitive relationship between *circ-MYOCD* and linear *MYOCD* splicing. SRPK3-mediated suppression of *circ-MYOCD* relieves the inhibition on linear splicing, leading to an accumulation of MYOCD protein. Given that MYOCD functions as a passive co-activator that forms concentration-dependent transcriptional condensates to activate cell identity genes [24], this “dosage” is critical. We propose that under physiological conditions, *circ-MYOCD* acts as a buffer to maintain MYOCD levels below the threshold required for pathological condensate formation. However, in the context of SRPK3 upregulation, the depletion of *circ-MYOCD* leads to excessive linear *MYOCD* production, likely facilitating aberrant transcriptional condensates that drive hyperactivation of downstream targets.

The role of MYOCD presents a therapeutic challenge due to its duality. While it drives pathological hypertrophy, it is also a cornerstone of cardiac regeneration (e.g., in the GMTM reprogramming cocktail) [25, 26]. Direct *MYOCD* ablation might impair reparative capacity. Targeting the upstream regulator SRPK3 offers a solution by restoring the natural feedback loop governed by *circ-MYOCD*. This approach “resets” the splicing balance, preventing pathological overexpression while likely preserving basal levels necessary for normal function.

In conclusion, we define the SRPK3-SRSF10-*circ-MYOCD* axis as a novel regulatory circuit that bridges splicing kinetics with circRNA-mediated signaling. By modulating the biophysical properties of nuclear MYOCD bodies and the composition of SRF transcriptional complexes, this axis dictates cardiac fate. Our findings underscore the “spliceosome-circRNA interactome” as a targetable frontier in heart failure therapy, supporting strategies such as pharmacological inhibition of SRPK3 or direct delivery of engineered *circ-MYOCD* mimics.

## Funding

This study was supported by the National Natural Science Foundation of China (No. 82170275, and 82170233).

## Author Contributions

Xiaowei Song, Jinchao Song, and Bi-Li Zhang designed the study. Xiaowei Song, Rui Cai, Feng Chen, Wenxia He, Zhenyu Zeng, Tianhong Ai and Shuting Hu performed experiments and data analysis. Xiaowei Song and Rui Cai prepared the figures and wrote and drafted the manuscript. All authors read and approved the manuscript.

## Competing Interests

All the authors have declared that no competing interest exists.

## Notes

### Competing Interest Statement

The authors have declared no competing interest.

